# Predicting Sex from Resting-State fMRI Across Multiple Independent Acquired Datasets

**DOI:** 10.1101/2020.08.20.259945

**Authors:** Obada Al Zoubi, Masaya Misaki, Aki Tsuchiyagaito, Vadim Zotev, Evan White, Tulsa 1000 Investigators, Martin Paulus, Jerzy Bodurka

## Abstract

Sex is an important biological variable often used in analyzing and describing the functional organization of the brain during cognitive and behavioral tasks. Several prior studies have shown that blood-oxygen-level-dependent (BOLD) functional MRI (fMRI) functional connectivity (FC) can be used to differentiate sex among individuals. Herein, we demonstrate that sex can be further classified with high accuracy using the intrinsic BOLD signal fluctuations from resting-state fMRI (rs-fMRI). We adopted the amplitude of low-frequency fluctuation (ALFF), and the fraction of ALFF (fALFF) features from the automated anatomical atlas (AAL) and Power’s functional atlas as an input to different machine learning (ML) methods. Using datasets from five independently acquired subject cohorts and with eight fMRI scanning sessions, we comprehensively assessed unbiased performance using nested-cross validation for within-sample and across sample accuracies. The results demonstrated high prediction accuracies for the Human Connectome Project (HCP) dataset (area under cure (AUC) > 0.89). The yielded accuracies suggest that sex difference is embodied and well-pronounced in the low-frequency BOLD signal fluctuation. The performance degrades with the heterogeneity of the cohort and suggests that other factors,.e.g. psychiatric disorders and demographics influences the BOLD signal and may interact with the classification of sex. In addition, the results revealed high learning generalizability with the HCP scan, but not across different datasets. The intraclass correlation coefficient (ICC) across HCP scans showed moderate**-**to-good reliability based on atlas selection (ICC = 0.65 [0.63-0.67] and ICC= 0.78 [0.76-0.80].). We also assessed the effect of scan duration on the predictability of sex and showed that sex differences could be detected even with a short rs-fMRI scan (e.g., 2 minutes). Moreover, we provided statistical maps of the brain regions differentially recruited by or predicting sex using Shapely values and determined an overlap with previous reports of brain response due to sex differences. Altogether, our analysis suggests that sex differences are well-pronounced in rs-fMRI and should be considered seriously in any study design, analysis, or interpretation.

## I. INTRODUCTION

Resting-state functional magnetic resonance imaging (rs-fMRI) is a non-invasive approach allowing for studies of brain functions by measuring hemodynamic flow within the resting brain. rs-fMRI has been proven to be an effective approach to discovering and studying consistent brain functional network organization (Biswal, 2012; Damoiseaux et al., 2006; Mantini, Perrucci, Del Gratta, Romani, & Corbetta, 2007). In particular, rs-fMRI has been used to identify differences across subjects based on demographic data and biological factors, including gender. Many studies have identified differences between males and females in terms of cognitive performance (Miller & Halpern, 2014), but these results do not provide a comprehensive and consistent view of sex differences (Del Giudice, 2009; Hyde & Plant, 1995). While there has been evidence of sex differences in some cognitive processes like language and emotional processing (Besson, Magne, & Schön, 2002; Schirmer, Kotz, & Friederici, 2005; Schirmer, Striano, & Friederici, 2005), other works could not found any conclusive evidence of such differences (Russell, Tchanturia, Rahman, & Schmidt, 2007; Wallentin, 2009). Similar to functional organization, sex differences were found in the structural organization of the brain (Chekroud, Ward, Rosenberg, & Holmes, 2016; Del Giudice et al., 2016; Rosenblatt, 2016). Research has shown that males have larger total brain volume, gray matter, and white matter tissues (Ingalhalikar et al., 2014). Also, intra- and inter-hemispheric connections have been shown to vary between males and females with a tendency for males to have higher intra-hemispheric connectivity (Ingalhalikar et al., 2014). In contrast, females showed high inter-hemispheric connectivity (Ingalhalikar et al., 2014). Moreover, brain regions like the insula, amygdala, and hippocampus have also been shown to structurally differ based on sex (Ruigrok et al., 2014). Similarly, authors in (Liu, Seidlitz, Blumenthal, Clasen, & Raznahan, 2020) reported consistent sex differences of gray matter volume (GMV) in the cortex and subcortical foci, brain regions associated with social and reproductive behaviors. This study also demonstrated a strong spatial coupling between brain regions showing GMV differences and brain expression of sex chromosome genes in adulthood. Despite the evidence of the brain structural differences, others have argued that both brain and behavior sex differences can be described as a mosaic of male and female properties with no clear binary distinction (Joel et al., 2015; Joel & Fausto-Sterling, 2016). Similarly, fMRI functional connectivity (FC) has been widely used to study sex differences. For instance, (Bluhm et al., 2008) reported an overall higher FC within the Default Mode Network (DMN) in the medial prefrontal and posterior cingulate cortices in females. Other work showed stronger inter-network FC in males and stronger intra-network FC in females (Allen et al., 2011). While there is much other evidence about sex differences in the resting state connectivity (Biswal et al., 2010; Tian, Wang, Yan, & He, 2011; Zuo, Kelly, et al., 2010), other works did not replicate nor consistently find any sex effects (Weis, Hodgetts, & Hausmann, 2019; Weissman-Fogel, Moayedi, Taylor, Pope, & Davis, 2010). Thus, investigating sex differences at the level of BOLD fluctuation may reveal if there is strong evidence of sex differences. Recently, machine learning (ML) techniques have been used widely to perform classification and regression on neuroscience data (Al Zoubi, Awad, & Kasabov, 2018; Al Zoubi, Ki Wong, et al., 2018; Campbell et al., 2020; Cohen, Chen, Parker Jones, Niu, & Wang, 2020; Du, Fu, & Calhoun, 2018; Garner et al., 2019; Kazeminejad & Sotero, 2019; Saccà et al., 2019). Some works focused on using ML for classifying subjects into male and female using functional (Ktena et al., 2018; Smith et al., 2013; Zhang, Dougherty, Baum, White, & Michael, 2018) and structural data (Chekroud et al., 2016; Feis, Brodersen, von Cramon, Luders, & Tittgemeyer, 2013; Rosenblatt, 2016). In this work, we focused on investigating sex classification using BOLD fMRI signal fluctuations. More specifically, BOLD can be characterized by the amplitude of the low-frequency fluctuation (ALFF) (Yu-Feng et al., 2007), which measures the extent of spontaneous fluctuation of the BOLD signal. ALFF has been linked to low-frequency oscillations from spontaneous neuronal activity and may manifest in the rhythmic activity and interaction of processing information across the brain (Cordes et al., 2001). ALFF is calculated by computing the power of the signal within [0.01-0.08] Hz or[0.01-0.1] Hz ranges (Li et al., 2017; Yu-Feng et al., 2007). In addition, other information can be derived from BOLD fluctuation like the fraction of ALFF (fALFF), which is defined as the ratio of the power within [0.01-0.08] Hz or [0.01-0.1] Hz ranges (Li et al., 2017; Q.-H. Zou et al., 2008) to the entire power within [0-0.25] Hz range. ALFF and fALFF have been used before to understand how intrinsic resting-state activity interacts at during cognitive task and resting-state activity(Fox, Snyder, Vincent, & Raichle, 2007; Mennes et al., 2011; Q. Zou et al., 2013). Furthermore, ALFF has been used to study different mental illnesses like schizophrenia (Alonso-Solís et al., 2017; Hoptman et al., 2010), attention deficit hyperactivity disorder(ADHD) (Yu-Feng et al., 2007), acute mild traumatic brain injury (Zhan et al., 2016), mild cognitive impairment (Bai et al., 2011) major depressive disorder (Wang et al., 2012) and many others.

Here, we provide comprehensive analyses on resting-state fMRI data, independently acquired from multiple large cohorts of individuals, to evaluate sex classification based on ALFF and fALFF features. To avoid the dimensionality problem (e.g., voxels from the whole brain vs. localized locations), we extracted ALFF and fALFF features averaged in the automated anatomical labeling atlas (AAL) (Tzourio-Mazoyer et al., 2002) and Power’s functional atlas (Power et al., 2011). We systematically compared various ML methods approaches for assessing sex classification for within-sample and across samples accuracies. We utilized a nested-cross-validation approach to avoid biased results that may arise from the use of traditional cross-validation. We studied the feasibility of deploying deep learning (DL) for sex classification in extension for emerged evidence of the utility of DL to analyze neuroscience data (He et al., 2020; Nguyen, Sun, Alexander, Feng, & Yeo, 2018; Pereira, Pinto, Alves, & Silva, 2016; Plis et al., 2014; van der Burgh et al., 2017; Vieira, Pinaya, & Mechelli, 2017). We assessed the importance of each feature using Shapley values (Lundberg & Lee, 2017) from both atlases. Then, we mapped the feature importance on the brain along with the direction of prediction. Recently, concerns about the test-retest reliability of rs-fMRI were raised (Noble, Scheinost, & Constable, 2019; Noble et al., 2017). Unlike the FC measures, ALFF has been shown to be reliable and reproducible across sessions (Zuo, Di Martino, et al., 2010). Thus, we examined the test-rest reliability of sex classification by calculating the Intraclass Correlation Coefficient (ICC) of sex classification from the HCP dataset. The effect of scan duration on sex classification was also assessed for the HCP dataset. Finally, the results from our comprehensive analyses will be discussed and summarized. The analyses offered here will allow to quantify the differences between males and females and evaluate the effect of psychiatric disorders on the ALFF and fALFF from the perspective of sex.

## II. Methods

### A. Datasets

Five datasets were used in this work to assess sex classification:

1. **ABIDE** Autism Brain Imaging Data Exchange database investigates the neural basis of autism (Di Martino et al., 2014). The data was collected from 16 international imaging sites and composed of 539 individuals suffering from autism spectrum disorders (ASD) and 573 typical controls. The data were preprocessed using the neuroimaging analysis kit (NIAK) pipeline described (Bellec et al., 2012), and only subjects with good data were used in this work. It should be noted that scan parameters, including the number of volumes, fMRI sequence repetition time (TR), and MRI scanners were different across the sites of data collection.
2. **HCP** Human Connectome Project dataset (S1200 release) is comprised of imaging data, including resting-state fMRI, from a large population of healthy young adults (Van Essen et al., 2013; Van Essen et al., 2012). We included the data from two rsfMRI sessions obtained over the course of two days. Each session consists of two scans with left-to-right (LR) and right-to-left (RL) phase encoding. We refer to the four scans as Ses11-RL, Ses1-LR, Ses2-RL, and Ses2-LR, respectively. The scan parameters were TR=720 ms, TE=33.1 ms and the number of volumes =1200. It should be noted that data were recorded using a multiband echo-planar imaging pulse sequence allowing for the simultaneous acquisition of multiple slices (Xu et al., 2013).
3. **ACPI**: Addiction Connectome Preprocessed Initiative dataset assess the effect of using cannabis on children diagnosed with ADHD. The readily preprocessed subjects were available through a Multimodal treatment study of ADHD (MTA). Scan parameters were TR=2170 ms, TE=4.33 ms, and the number of volumes=180.
4. **COBRE-MIND**: Center for Biomedical Research Excellence – Multimodal Neuroimaging of Neuropsychiatric Disorders (Calhoun et al., 2012; Mayer et al., 2013) dataset from 72 patients with schizophrenia and 75 healthy controls. Preprocessed subjects were available through the NIAK preprocessing pipeline. Scan parameters were TR=2000 ms, TE=29 ms, and the number of volumes =150.
5. **T1000**: We used the first 500 subjects of the Tulsa 1000 (T-1000), a naturalistic study assessing and longitudinally following 1000 individuals, including healthy individuals and treatment-seeking individuals with substance use, eating disorders, and mood disorders and/or anxiety (Victor et al., 2018). Scan parameters: TR=2000, TE=27 ms, and a number of volumes =240.

In addition, Table 1 shows the final number of samples and the sex distribution across the five datasets.

**Table 1.** Datasets used to predict sex from fMRI resting scans.

| Dataset | # Samples | Female/Male | Population |
| --- | --- | --- | --- |
| T1000 | 426 | 272/154 | Healthy controls, mood/anxiety, substance use, and eating disorders |
| HCP-Session1-RL | 1047 | 566/481 | Healthy young subjects |
| HCP-Session1-LR | 1065 | 573/492 | Healthy young subjects |
| HCP-Session2-RL | 1004 | 537/467 | Healthy young subjects |
| HCP-Session2-LR | 987 | 526/461 | Healthy young subjects |
| ABIDE | 871 | 144/727 | Healthy control and autism spectrum disorders |
| COBRE-MIND | 146 | 37/109 | Schizophrenia and healthy controls |
| ACPI | 126 | 25/101 | Substance use and ADHD |

### B. Preprocessing pipelines

We relied on publicly available preprocessed datasets, if existing, to avoid any biases that could arise from reprocessing data. If possible, we tried to match the preprocessing pipelines as well. For ABIDE and COBRE-MIND datasets, the preprocessed data were obtained through the NIAK pipeline (Bellec et al., 2012) without global signal regression options. For ACPI, the preprocessed data were available through the configurable pipeline for the analysis of the connectomes (C-PAC) pipeline (Craddock et al., 2013; Lurie et al., 2013) without motion scrubbing and no global signal regression. For HCP, we used ICA-FIX denoised rs-fMRI volumetric data available in (HCP S1200 release). The data were spatially normalized to MNI152 at the time of download. We did not apply any additional noise corrections to the data. Subjects with relative Root Mean Square (RMS) motion>0.2 were further excluded. Finally, for T1000, we applied the following preprocessing steps, including despike, cardiac- and respiration-induced noise reduction RETROICOR (Glover, Li, & Ress, 2000), Linear warping to the MNI space. We also applied another layer of noise reduction by regressing out low-frequency, 12 motion parameters, local white matter average signal (ANATICOR) (Jo, Saad, Simmons, Milbury, & Cox, 2010), and three principal components of the ventricle signal from the signal time course. As mentioned above, subjects with RMS motion larger than 0.2 were also excluded from the analysis.

### C. ROI definition

In this analysis, we relied on predefined anatomical and functional atlases. First, we used Automated Anatomical Labeling (AAL) (Tzourio-Mazoyer et al., 2002), which includes 116 ROIs that expand across the whole brain. On the other hand, we used Power’s ROIs compromised of 264 ROIs (Power et al., 2011). For each atlas, we extracted the average time series from ROI voxels after detrending the signal.

### D. ROI-based Features

The ALFF was computed as the signal power within 0.01 and 0.1 Hz range of the average time series of each ROI. The fALFF was calculated as the ratio of signal power within 0.01 to 0.1 Hz range to the total power within 0 and 0.25 Hz range. This resulted in 264 (ALFF, fALFF) pair of values for Power ROIs atlas and 116 (ALFF, fALFF) pair values for AAL atlas.

### E. Machine Learning Method

#### Classical Machine Learning (ML) methods

We considered several ML methods, including support vector classification (svc) with both linear and radial basis function (RBF), Random Forests Classifier (RandomF), logistic regression with *ℓ1-norm* (logistic_l1) or *ℓ2-norm* (logisitc_l2), gaussian naive bayes (GaussianNB) and extreme gradient boosting (xgboost) algorithm (Chen & Guestrin, 2016). The Scikit-learn machine learning package (Pedregosa et al., 2011) was used to implement each classifier.

#### Deep Learning (DL) methods

We adopted three models to obtain spatial information from the rsfMRI feature. The architectures deployed one-dimensional convolution (Conv 1D) layers while treating each sub-feature as a different channel. We used kernel size of *k=3*, stride *s=1* and filter size with an order of *f=16*. The activation was set ‘ReLU’ function. In addition, we used Max Pooling layers before dropout layers (p=0.4) to improve the generalizability of the DL models. The models were generated by increasing the number of blocks from 1 to 3. Each time we added a new block, we increased the number of by 16×N with N as the block number. We also increased the number of neurons in the fully-connected layer based on the number of added blocks to have 100, 200, and 400, respectively. TensorFlow with Keras backend was used to build and train the three models. Adam optimizer with early stopping callback (patience =10, validation =30%) was utilized after setting the maximum number of epochs to 500 and batch size =64.

### F. Evaluation strategy

We utilized the area under the curve (AUC) for reporting the results. AUC is less sensitive for imbalanced classes and offers a robust measure for binary classification problems (Ling, Huang, & Zhang, 2003). Using 10 repeats, the average AUC is adopted to select the best classifier or atlas. In addition, we used stratified nested-cross-validation (sNCV) instead of the classical cross-validation to preserve the ratio of samples between the groups while providing unbiased estimation. The sNCV avoids biased results by isolating testing data from any parameter optimization. We used an inner loop of three-fold cross-validation to optimize each classifier’s parameters. Then, the model with the best performance was used to extract the prediction from the testing set. We always report the AUC for the testing set and refer to it as an out-of-sample performance. We followed three evaluation strategies to evaluate sex classification. First, we assessed the performance of each ML approach on each dataset (within-sample evaluation). Secondly, we used leave-one-scan-out (across samples evaluation) to test the reproducibility of predicting sex across different datasets. Thirdly, we focused on HCP to evaluate the effect of scanning time on the predictability of sex. More specifically, we varied the number of samples [32, 150, 300, 600, 900, 1200] and calculated the AUC accordingly. The analysis was applied to both atlases for only the best classifiers found from previous analyses. Finally, we investigated the ICC from HCP to evaluate the consistency of predictions across HCP scans. ICC measures the amount of variability that can be explained by an objective of measurement, such as subject (Shrout & Fleiss, 1979). We reported the ICC (2,1), which is used to estimate the agreements (predicted probabilities) when the sources of error are known (multiple scans from HCP). To offer an accurate estimation of ICC, we split each scan’s data into 50-fold sNCV and estimated the probability of each sample in the testing set. We repeated the probability estimation for all scans using AAL and Power’s atlases. It should be noted that only the best ML method found from previous analyses was used to estimate the predicted probability of sex.

### G. Feature importance

In order to reveal and map the important features for sex classification, while providing interpretable results, we propose using the Shapley Additive (SHAP) approach (Lundberg & Lee, 2017; Štrumbelj & Kononenko, 2014). SHAP deploys a game-theoretical approach to estimate Shapley Values (SHV) of a cooperative game while assuming each feature as an independent player. To compute SHV, each feature goes under random sampling and substantiation to assess the impact of those features on the overall prediction. In our analysis, we used the best classifier obtained in the analysis as an Explainer. The process was done in 10-fold-cross validation with five repeats. The final SHV values were obtained as the average of out-of-sample prediction within each scan. The sign and strength of SHV values represent the importance of predicting and direction of prediction (positive is males, and negative is females based on our class encoding).

## III. Results

First, we investigated the performance of the classical ML and DL in predicting sex across all the five datasets. Figure 2 shows a boxplot of the area under the curve AUCs classification performance using classical ML and DL models (10 repeats with 10-fold sNCV).

**Figure 1.**
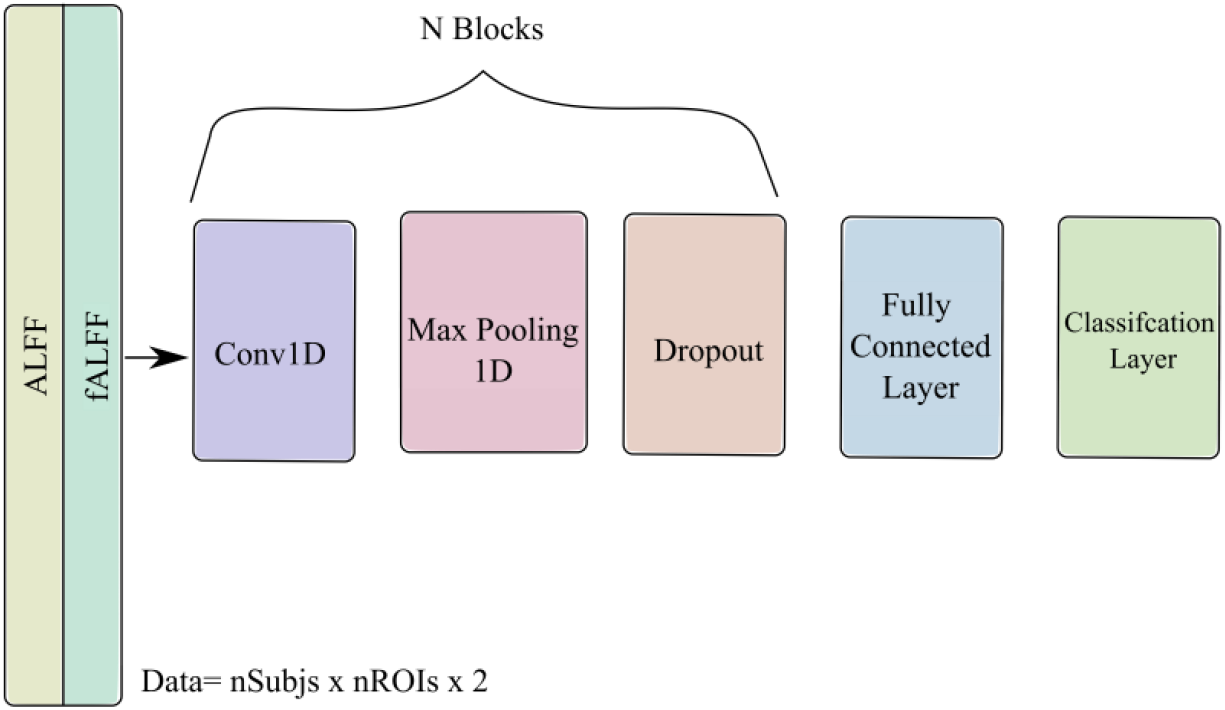
Deep Learning architecture for sex classification. The architecture consists of N-block of stacked of a convolutional layer, max pooling, and dropout layers. The previous architecture resulted in three models based on N=1,2 and 3.

**Figure 2.**
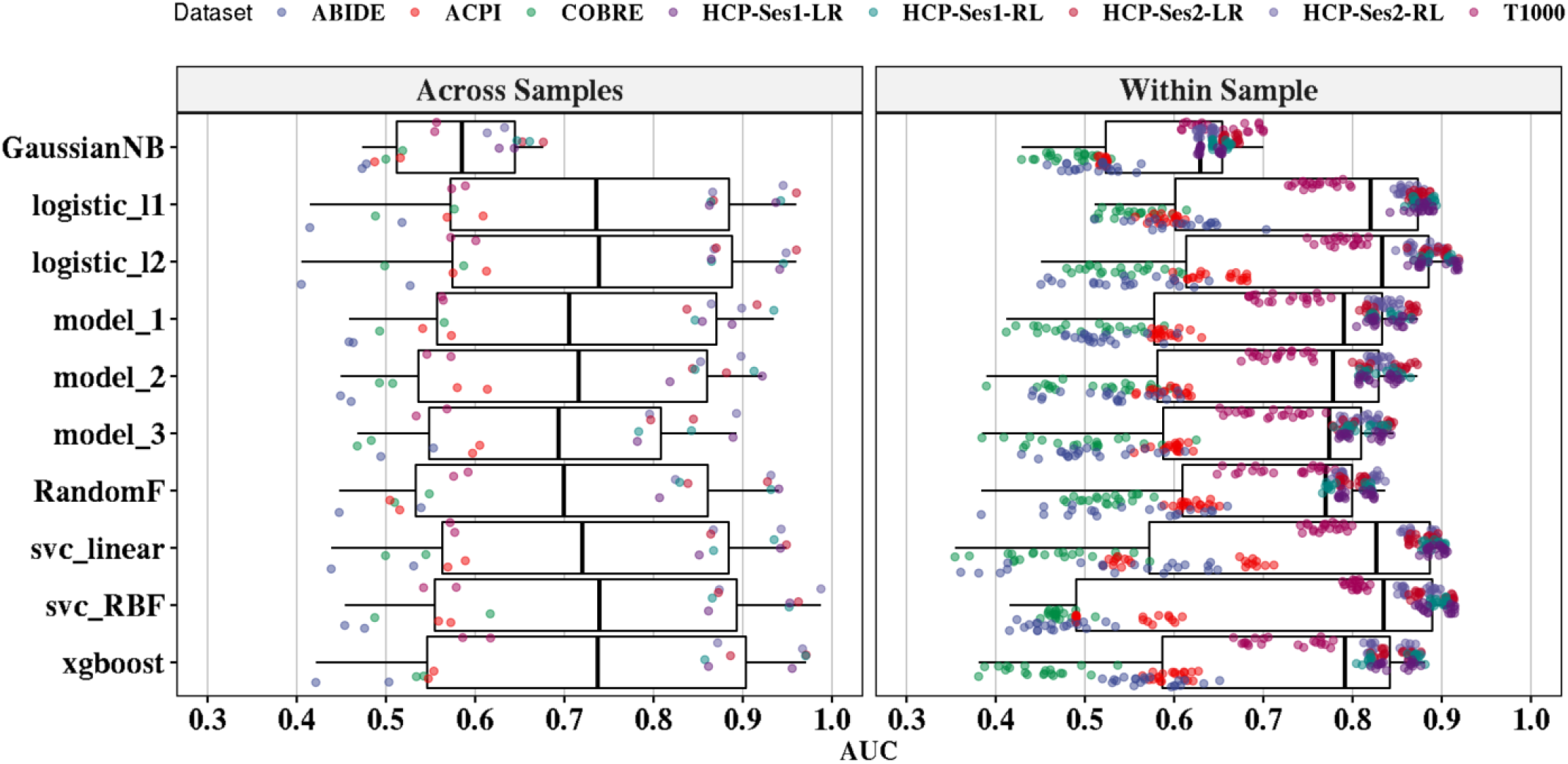
Binary classification performance of individual classifiers from classical ML and DL models. Performance is reported across and within-sample validation. Across samples (left panel), each point represents the AUC value of the leave-one-scan-out for each classifier using each atlas and each scan (2 atlases × 8 scans). For within-sample (right panel), each point is the out-of-sample AUCs values after running classifiers on each atlas and each scan with repeats (10 repeates×10 folds × 2 atlases × 8 scans).

Logistic regression methods with *ℓ2-norm* and *ℓ1-norm* yielded, on average, the best AUCs of 76% (±15%) 74.1% (±13.7%), respectively, for within-sample classification. Other classifiers achieved the following accuracies: svc with linear kernel [74.1% (±17.3)], svc with RBF kernel [72.9% (±19)], xgboost [71.6% (±15.1)], random forest [70.2% (±11.8)] and Gaussian Naive Bayes [59.5% (±7.3)]. Similarly, logistic regression with *ℓ2-norm* achieved the best accuracy of 72.8% (±19.4) for Across sample evaluation, followed by xgboost. Thus, we selected and used logistic regression with *ℓ2-norm* for our further analysis.

In Figure 3, we show the performance of logistic regression with *ℓ2-norm* performance on the individual datasets using across and within-sample evaluation.

**Figure 3.**
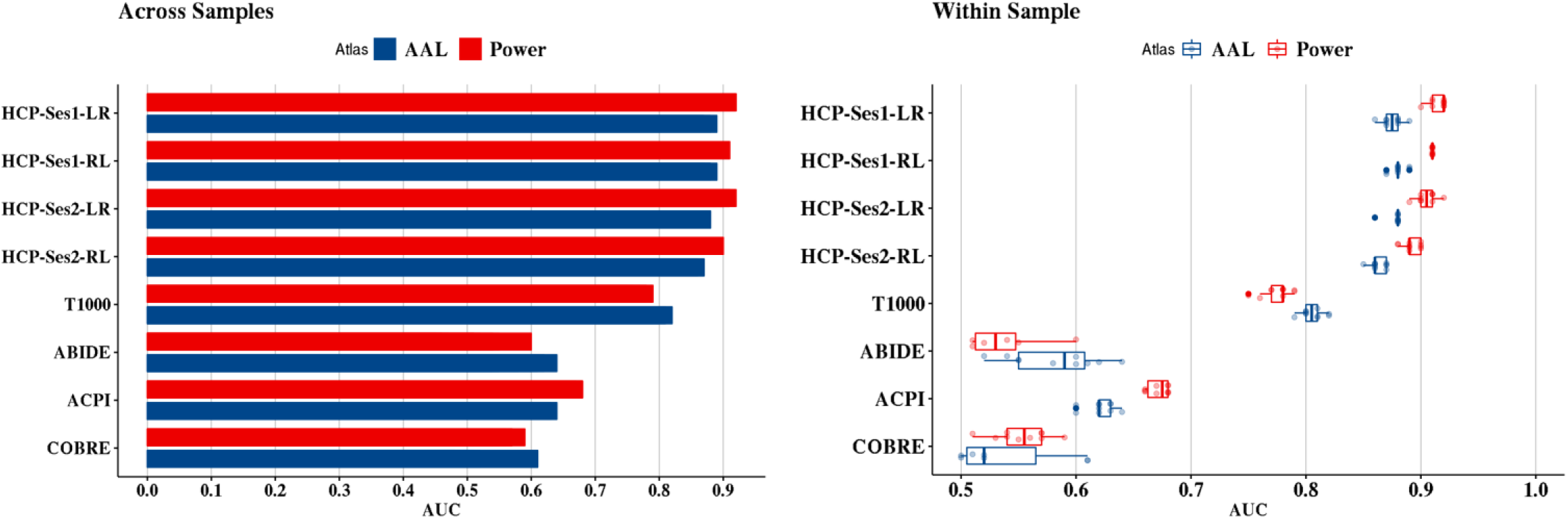
The yielded out-of-sample AUC values using logistic regression with ℓ2-norm based on each atlas. The left panel represents the across samples evaluation, while the right panel represents the within-sample evaluation from each of the 10 repeats.

We also compared the performance of classifiers based on the adopted atlas (Figure 4). From the analysis, Power’s functional atlas achieved the highest average AUC of 71.8% (±20) and 71.6% (±15.6) for across and within-sample evaluation. AAL achieved 68.2% (15.7) for across sample evaluation and 70.6% (±14.5) for within-sample evaluation.

**Figure 4.**
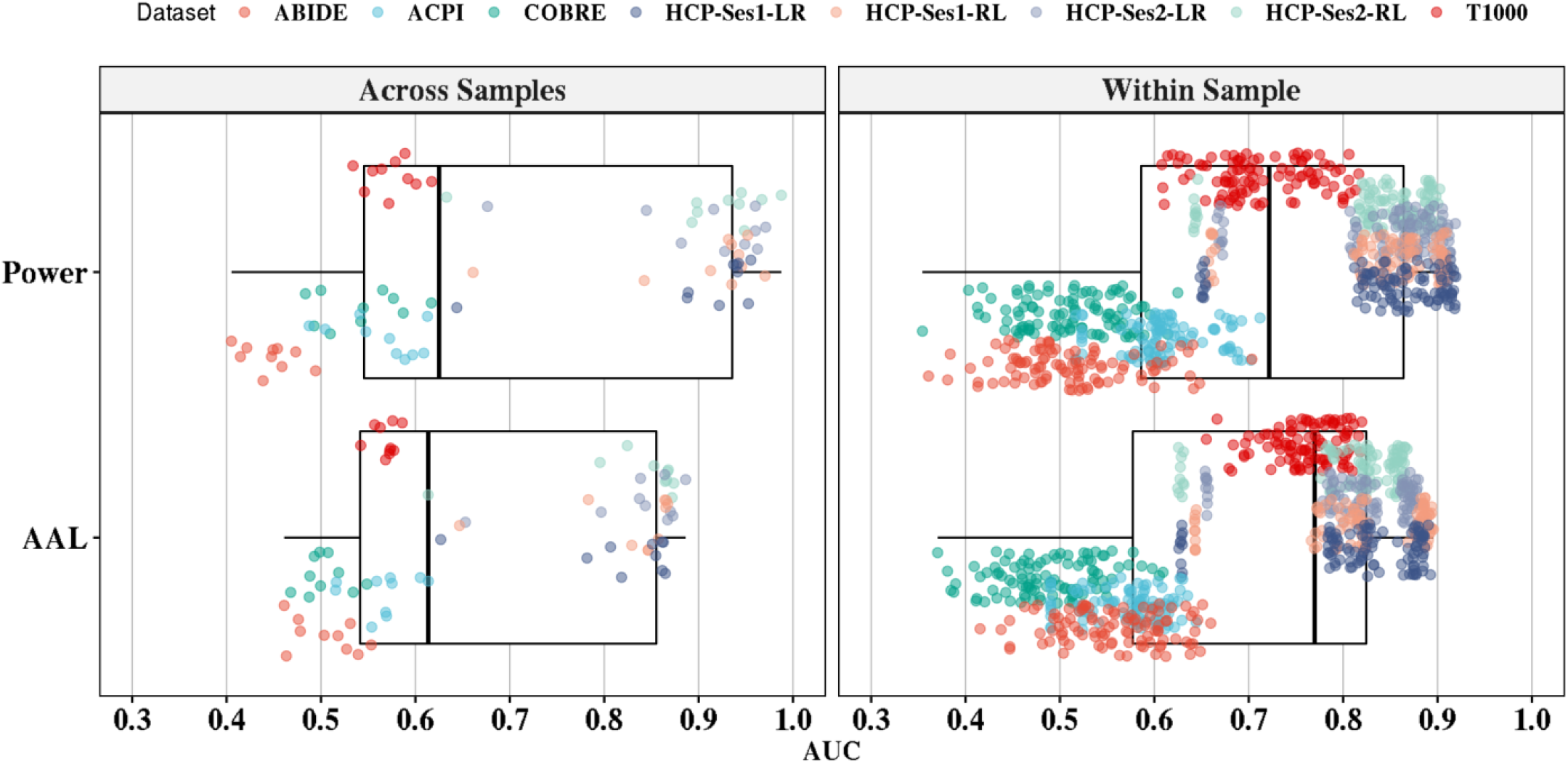
The effect of atlas selection of the performance of sex classification. The left panel represents the out-of-sample AUC values, averaged across all classifiers (10 classifiers × 8 scans). The right panel represents the within-sample AUC values across all classifiers (10 repeates×10 folds × 10 classifiers × 8 scans).

The effect of the number of samples on the prediction accuracies is shown in Figure 5. Power’s functional atlas performed better than AAL when fixing the number of samples. The accuracy does not seem to improve after 600 to 900 time points for Power’s atlas; however, the AAL improves over the number of samples but does not reach Power’s atlas performance.

**Figure 5.**
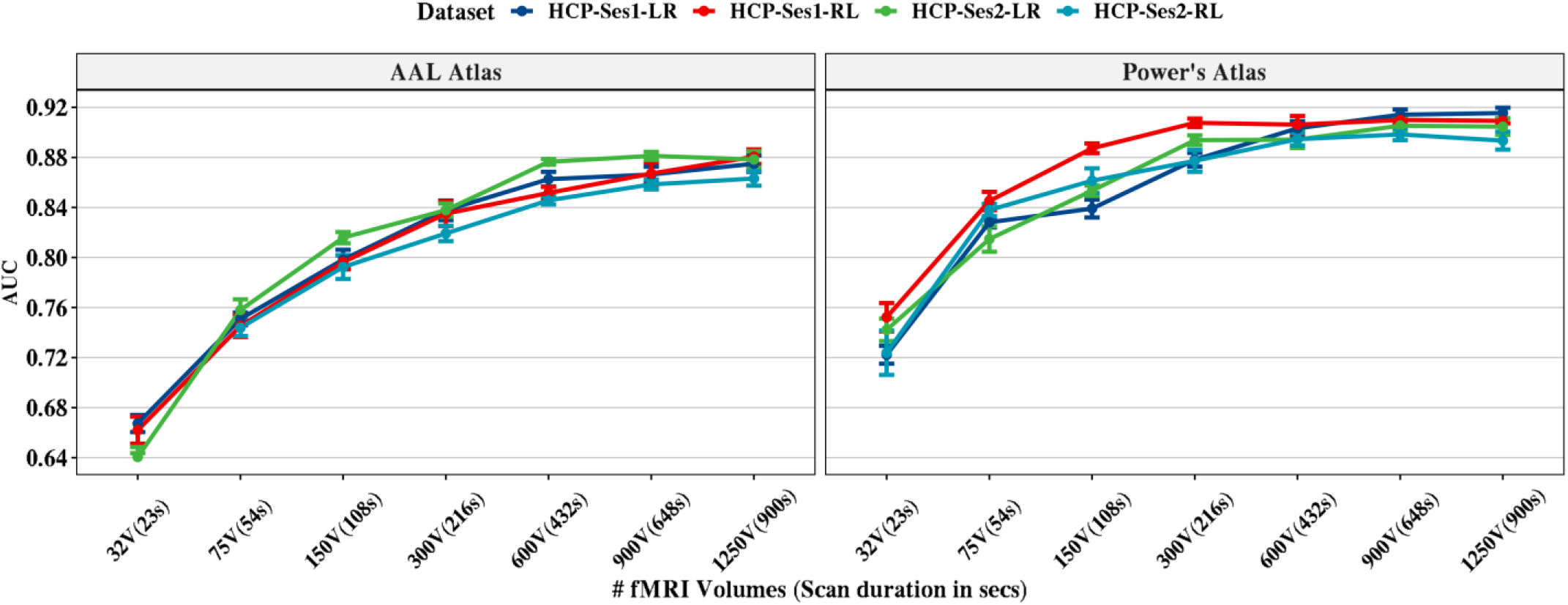
The effect of the number of volumes on sex classification from HCP scans. For each of the number of volumes, we ran 10-repeat of 10-fold sNCV on the data and reported the out-of-sample AUC values using logistic regression with the ℓ2-norm classifier. The left panel represents the AAL atlas performance, and the right panel shows the Power’s atlas performance. The error bars represent the standard deviation of AUCs from the 10-repeats (10 repeates×10 folds).

We also assessed the test-retest reliability by calculating the ICC using the four scans of the HCP dataset. We used the logistic regression with the ℓ2-norm to estimate the predicted probabilities of sex with sNCV configuration (testing set). As in the previous analyses, we used combined ALFF and fALFF features as an input for the logistic regression with the ℓ2-norm. The results indicated moderate reliability for AAL with ICC=0.65 [0.63-0.67] and good reliability for Power’s atlas with ICC=0.78 [0.76-0.80].

Feature importance for AAL and Power’s atlases is shown in Figure 6 and Figure 7. For Power’s ROIs, the size and color of the nodes represent the importance of those nodes in predicting sex from the five datasets. The importance was computed using SHAP values from logistic regression with the *ℓ2-norm* explainer. The red color represents the importance of predicting females, while blue represents predicting males. Similarly, we mapped the SHAP values for AAL on the surface of the brain while using the same color-coding used in Power’s atlas. It should be noted that the SHAP values were calculated for the out-of-sample prediction and averaged over 5-repeats.

**Figure 6.**
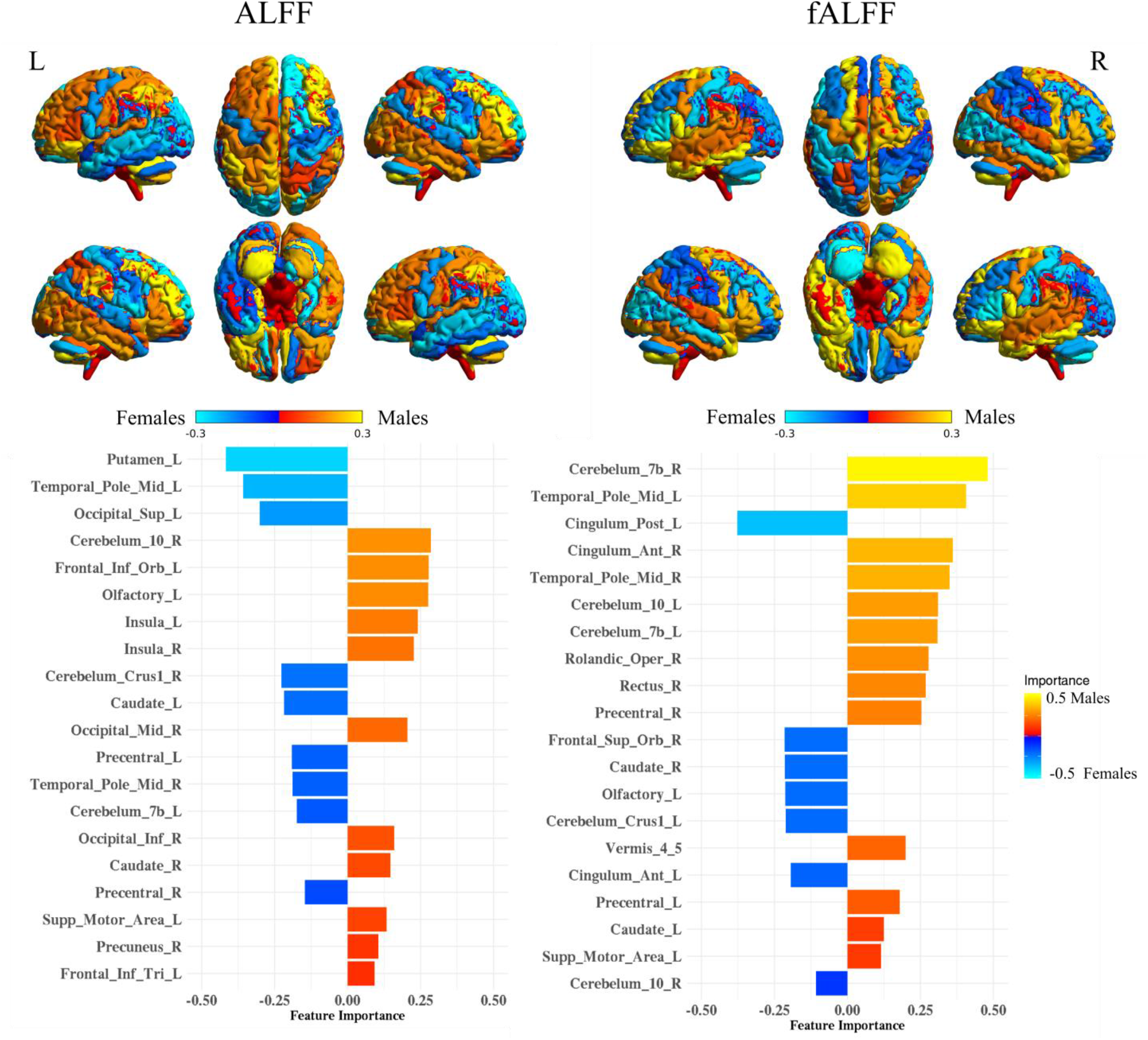
ALFF and fALFF feature maps and importance in sex classification using SHAP values (SHV) for AAL atlas. The colors are mapped based on the SHV values and reveal the contribution of each region in sex classification. The bar plot shows the top 20 features ordered based on the absolute SHV values.

**Figure 7.**
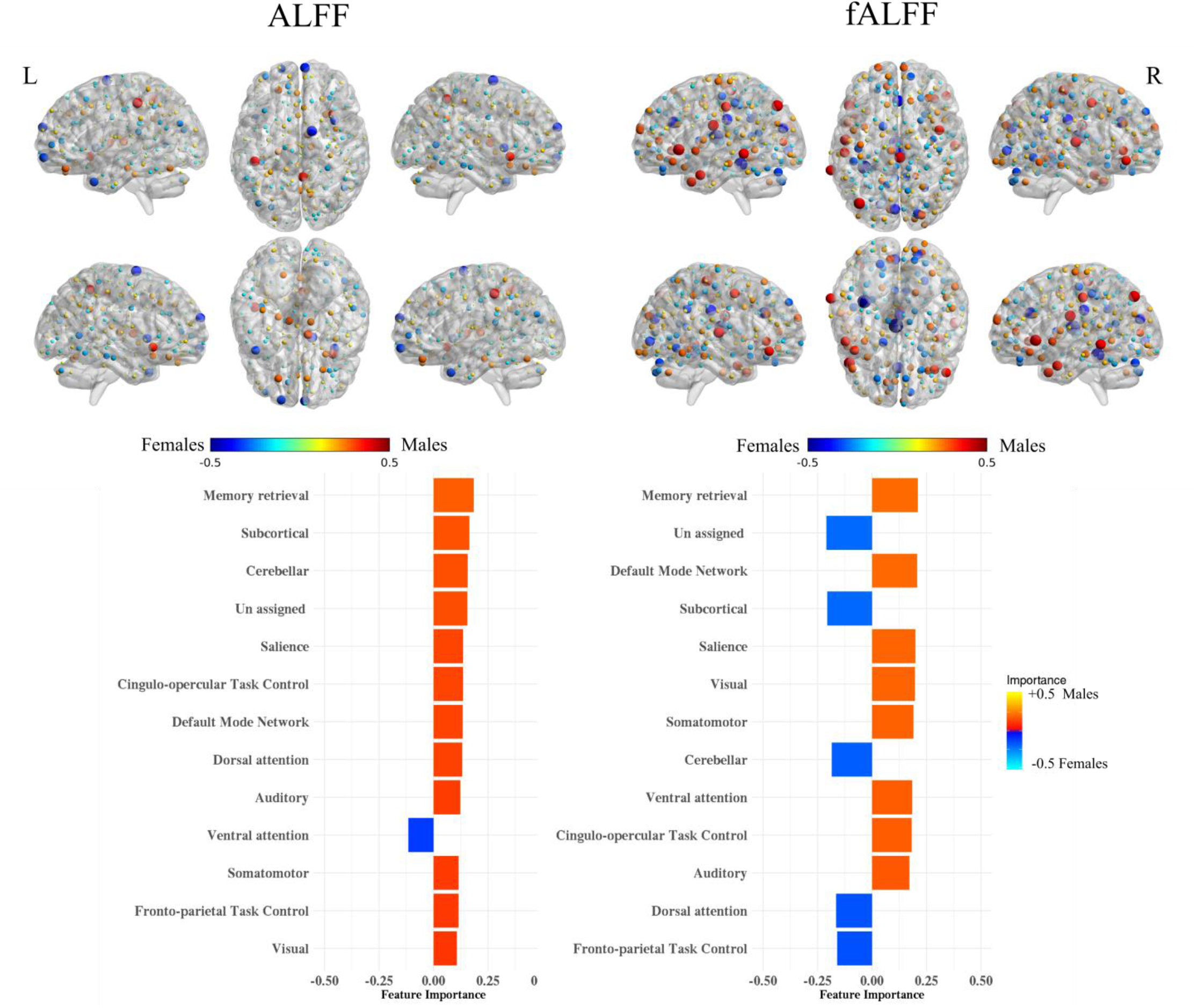
ALFF and fALFF feature maps and importance in sex classification using SHAP values (SHV) for Power’s functional atlas. The colors are mapped based on the SHAP values and reveal the contribution of sex classification. For the bar plot, the 264 regions were aggregated based on the assigned brain system (Power et al., 2011) and ordered based on the mean absolute SHV values.

## IV. Discussion

We conducted comprehensive analyses to predict sex from rsfMRI across five independent acquired datasets and structured discussion as follows.

### A. Predictability of Sex

We show that males and females can be classified with high accuracy in healthy young adults when using intrinsic BOLD fluctuation properties while deteriorating in heterogenous datasets (Figure 2). To avoid the “curse of dimensionality,” we focused on the region-of-interest (ROI) approach to characterize the BOLD fluctuation properties rather than using whole-brain data. This allowed us to have a robust prediction and avoid potential overfitting (Guyon & Elisseeff, 2003; Hua, Tembe, & Dougherty, 2009; Mwangi, Tian, & Soares, 2014). We derived ROIs of interests from two atlases, namely, Power’s functional atlas and AAL anatomical atlas. Both atlases are widely used in analyzing rsfMRI data and manifest different methodologies in parcellating the brain. While Power’s atlas uses the functional organizations of the brain, dividing it into 264 ROIs, AAL atlas relies on the anatomical distribution of the brain, categorizing it into 116 brain regions. Overall, we found that sex is predictable with the highest accuracy in healthy young adults (HCP dataset). The more heterogeneous the dataset becomes, including the mental illness factor, the less predictable the sex is. Other used datasets varied in population and mixed clinical symptoms, with the best sex prediction performance achieved in the T1000 dataset. Our findings support and extend the good sex classification results based on fMRI functional connectivity, as shown in (Weis et al., 2020; Zhang et al., 2018). Additionally, it supports the notation that mental illnesses disrupt the properties of BOLD fluctuation as it has been shown in several clinical populations like autism (Itahashi et al., 2015; Noonan, Haist, & Müller, 2009), ADHD (Tang, Wei, Zhao, & Nie, 2018), schizophrenia (Hoptman et al., 2010; Yu et al., 2014), bipolar disorder (Meda et al., 2015; Yang et al., 2019) and depression (Jing et al., 2013). Altogether, the results may suggest that sex difference is primarily encoded in the low-frequency BOLD fluctuations as characterized in the ALFF and fALFF measures and can be potentially used as a biomarker for analyzing different clinical populations.

### B. Effect of Classifier on Sex Predictability

We investigated the choice of the classifier on the performance of sex classification using several classical ML methods in addition to DL. The results revealed that linear classifiers outperformed both non-linear classifiers and DL models with the best average AUC value using logistic regression with *ℓ2-norm*, followed by logistic regression with *ℓ1-norm* regularization. The performance of classifiers was evaluated using across and within samples. The performance of predicting sex varied across the datasets and scans with the best performance using the four scans of HCP—the performance of classification degraded as a function of the heterogeneity of the sample. Thus, BOLD fluctuation properties may be largely impacted by the clinical diagnosis and can thus potentially be biomarkers for clinical symptoms.

### C. Effect of ROI Selection

Both atlases yielded close accuracies with an advantage for Power’s atlas. More specifically, Power’s atlas achieved higher average AUC than AAL atlas for all datasets except for T1000 and ABIDE datasets (using l2-nom logistic regression). Similarly, Power’s atlas resulted in better AUC for all datasets except for T1000, ABIDE, and COBRE-MIND datasets when using across sample evaluation. The difference in the performance may be attributed to the disease-specific alteration for brain structural and functional originations of the brain.

### D. Generalizability

We tested the generalizability of sex classification by using an across datasets evaluation approach; we trained on all scans except one, which was then used for testing. The analysis resulted in one AUC per dataset and atlas. We compared both classical ML and DL to investigate the generalizability of each classifier. The results indicated poor generalizability across datasets except for HCP. The fact that HCP is comprised of multiple scans recorded from the same subjects has contributed to high AUC within each scan. As in the within-sample evaluation, linear classifiers outperformed non-linear classifiers with the advantage of *ℓ2-norm* logistic regression over other classifiers. DL models did not generalize very well, yielding results similar to the non-linear classical ML methods. Thus, further research should be conducted in order to find suitable ML techniques for brain imaging data that account for variability across subjects, a limited number of samples, and high dimensional data.

### E. The effect of the number of samples

The effect of the number of samples (e.g., resting fMRI scan duration) on sex classification was evaluated on HCP scans since they have the longest scan time (∼ 15 minutes). For each scan, we took the first s= [32, 75, 150, 300, 600,900, 1200] samples and extracted ALFF and fALFF features. Each time, we accessed the AUC for within-sample 10-repeats of 10-fold NCV. Using only the first 32 samples from HCP scans, AUCs were between 0.66 to 0.72. The performance for Power’s atlas seems to plateau between 600 and 900 samples with little improvement after adding more samples. AAL atlas performance was lower than Power’s atlas for the same number of points. Thus, researchers should account for sex differences for experiments with even short innervation design (e.g., block design experiments).

### F. Test-Retest Reliability

The test-rest reliability of sex classification was assessed by calculating the ICCs from HCP scans. The results indicated moderate reliability for AAL with ICC=0.65 [0.63-0.67] and good reliability for Power’s atlas with ICC=0.78 [0.76-0.80]. The moderate and good reliability from the HCP dataset offers promising results for using ML to analyze rs-fMRI instead of the traditional FC analysis of rs-fMRI. The reliability of ALFF and fALFF has been shown to be reliable across sessions (Zuo, Di Martino, et al., 2010), unlike the reliability of the classical FC analysis of rs-fMRI (Noble et al., 2019; Noble et al., 2017), which lead many researchers to endorse the notion of the “reproducibility crisis” for FC (Baker, 2016). Thus, the reliability of low-frequency fluctuation properties across sessions, along with moderate to good prediction reliability, make them better measures to study and characterize brain functional responses in health and disease.

### G. Spatial distribution and feature importance

We adopted the SHAP approach to assessing feature importance and directionality in predicting each sex. For AAL atlas, we mapped the SHAP values on the surface of the brain. The results revealed that sex classification is not associated with one specific region, but varies across the brain and sub-features set. Also, there are no regions that are associated explicitly with differentiating females from males. However, some brain regions are consistently among the top important parts in predicting sex, like the cerebellum and temporal pole for ALFF and fALFF. Top features in our case span over part of the DMN, temporal pole, precuneus, and insular cortex regions. Additionally, we observed an overlap for top brain regions differentiating sex - in our case sex difference analysis using GVM analysis (Liu et al., 2020) and FC analysis (Weis et al., 2020). Also, we replicated the observation that the DMN is one of the top features in differentiating sex, in line with the findings from (Biswal et al., 2010; Zhang et al., 2018).

For the Power’s functional atlas, we plotted the top features using node size and color. Some ROIs overlap with top important features from the AAL, like in the DMN and temporal pole. We reported average SHAP values by averaging them by the brain system (Power et al., 2011) and showed that brain regions involved in memory retrieval constitute the top predicting features in both ALFF and fALFF features (Figure 7). Overall, the obtained distribution of feature importance supports the notion that the brain consists of mosaic features (Joel et al., 2015; Joel & Fausto-Sterling, 2016), where some features are more pronounced in one sex than in the other. In our case, the mosaic features are not only spatially distributed but also span across ALFF and fALFF features.

## V. Limitations

In this work, we explored using rs-fMRI to predict sex from five independent datasets. The datasets were collected at various sites using different MRI scanners, populations, preprocessing pipelines, and other configurations. The effect of these factors on predicting sex is still not apparent nor well characterized (Botvinik-Nezer et al., 2020). In addition, we used two AAL and Power’s atlas only. There are many other functional and anatomical atlases that can be adopted. We used several ML and DL methods, but there are many other ML methods and DL architectures that have not been explored here. We used ALFF and fALFF for describing BOLD signal fluctuations; however, other features can be defined and used.

## VI. Conclusion

Machine learning (ML) has gained popularity in predicting different outcomes in human brain neuroimaging data. In this work, we have shown that sex can be predicted with high accuracy from resting-state fMRI using various ML methods. The results demonstrated that sex difference is embedded in the properties of low-frequency BOLD signal fluctuation and extends the previous findings of sex difference reported based on fMRI functional connectivity. We assessed the sex classification on of five different and independent datasets that vary in population, including healthy young adults to other clinical populations. The highest archived results occurred when using healthy young adults only and may reflect the effect of the mental illnesses on the properties of the BOLD signal. The best classification performance was obtained with the use of linear classifiers, and we did not find an advantage of using Deep Learning methods. The spatial distribution of the important features was consistent with the previous finding, but we showed that sex classification did not rely on a specific brain region nor one sub-feature set. The results presented here suggest that sex distribution should be seriously considered in any brain imaging study including studies that investigate functional connectivity, BOLD activation or structural analyses.

## VII. Acknowledgment

This work has been supported directly by The William K. Warren Foundation, Laureate Institute for Brain Research, and in part by the P20 GM121312 award from the National Institute of General Medical Sciences, National Institutes of Health. We would also like to thank the open-source data community and preprocessed data initiatives for giving access to rest-fMRI datasets and funding agencies. The funding and support for each dataset are as follows: (1) ACPI is primarily supported by a grant supplement (R01 MH094639) by the National Institute on Drug Abuse (NIDA). Additional support provided by the Child Mind Institute and the Nathan Kline Institute; (2) ABIDE: the Autism Brain Imaging Data Exchange for preprocessing and providing the data; (3) COBRE-MIND: the National Institute of Health Center of Biomedical Research Excellence grant 1P20RR021938-01A2; (4) HCP: data were provided [in part] by the Human Connectome Project, WU-Minn Consortium (Principal Investigators: David Van Essen and Kamil Ugurbil; 1U54MH091657) funded by the 16 NIH Institutes and Centers that support the NIH Blueprint for Neuroscience Research; and by the McDonnell Center for Systems Neuroscience at Washington University.”

